# Cirscan: a shiny application to identify differentially active sponge mechanisms and visualize circRNA-miRNA-mRNA networks

**DOI:** 10.1101/2023.11.07.564256

**Authors:** Rose-Marie Fraboulet, Yanis Si Ahmed, Marc Aubry, Sebastien Corre, Marie-Dominique Galibert, Yuna Blum

**Affiliations:** Univ Rennes, CNRS, INSERM, IGDR (Institut de Genetique et Developpement de Rennes) - UMR 6290, ERL U1305, F-35000 Rennes, France; INSERM, OSS (Oncogenesis Stress Signaling), UMR S 1242, CLCC Eugene Marquis, Univ Rennes 1, 35000, Rennes, France; Department of Molecular Genetics and Genomics, Hospital University of Rennes (CHU Rennes), F-35000 Rennes, France

**Keywords:** Circular RNAs, sponge mechanism, transcriptomic data, regulation network, cancer

## Abstract

**Background:** Non-coding RNAs represent a large part of the human transcriptome and have been shown to play an important role in disease such as cancer. However, their biological functions are still incompletely understood. Among noncoding RNAs, circular RNAs (circRNAs) have recently been identified for their microRNA (miRNA) sponge function which allows them to modulate the expression of miRNA target genes by taking on the role of competitive endogenous RNAs (ce-circRNAs). Today, most computational tools are not adapted to the search for ce-circRNAs or have not been developed for the search for ce-circRNAs from user’s transcriptomic data.

**Results:** In this study, we present Cirscan (CIRcular RNA Sponge CANdidates), an interactive Shiny application that automatically infers circRNA-miRNA-mRNA networks from human multi-level transcript expression data from two biological conditions (e.g. tumor versus normal conditions in the case of cancer study) in order to identify on a large scale, potential sponge mechanisms active in a specific condition. Cirscan ranks each circRNA-miRNA-mRNA subnetwork according to a sponge score that integrates multiple criteria based on interaction reliability and expression level. Finally, the top ranked sponge mechanisms can be visualized as networks and an enrichment analysis is performed to help its biological interpretation. We showed on two real case studies that Cirscan is capable of retrieving sponge mechanisms previously described, as well as identifying potential novel circRNA sponge candidates.

**Conclusions:** Cirscan can be considered as a companion tool for biologists, facilitating their ability to prioritize sponge mechanisms for experimental validations and identifying potential therapeutic targets. Cirscan is implemented in R, released under the license GPL-3 and accessible on GitLab (https://gitlab.com/geobioinfo/cirscan_Rshiny). The scripts used in this paper are also provided on Gitlab (https://gitlab.com/geobioinfo/cirscan_paper).

## Background

Non-coding RNAs (ncRNAs) are by definition, genome sequences not translated into proteins. For years, their biological functions was largely underestimated as they were considered to be junk transcriptional products. Recently, it has however been demonstrated that ncRNAs represent a large part of the human transcriptome and have important functional roles, particularly in cancers [1]. Advances in sequencing technologies have led to the identification of different types of ncRNAs in the cell, such as long non-coding RNAs and microRNAs (miRNAs) [2]. In this study, we are interested in circular RNAs (circRNAs), a category of recently discovered ncRNAs known for their miRNA sponge function [3, 4]. MicroRNAs typically bind to mRNA through miRNA Recognition Elements (MREs), resulting in the degradation of their mRNA targets. However, when a circRNA shares MRE sites for a specific miRNA, it can function as a competitive endogenous RNA (ce-circRNA): the circRNA acts as a sponge, sequestering the miRNA and preventing it from binding to its mRNA targets, thereby indirectly regulating the mRNA expression [5, 6, 7]. One of the most wellknown ce-circRNA is ciRS-7 (CDR1as), mainly expressed in the brain, which acts as a regulator of the miR-7 with more than 70 binding sites and leads to increased expression levels of miRNA-7 targets [8, 9]. Thanks to their closed-loop structure formed by reverse-splicing, circRNAs are very stable and their capacity to bind to miRNAs seems to be stronger than that of any other ceRNA, leading to their super-sponge naming [10]. These interactions between coding and non-coding RNAs can be represented as circRNA-miRNA-mRNA interaction networks.

Few computational tools have been developed for the identification of ceRNAs but are not adapted to the identification of ce-circRNAs. This constraint is primarily due to their input requirements, which accept only two types of RNA dataset simultaneously (i.e. miRNA and mRNA expression data), and to their predictions databases restricted to interactions between miRNA and mRNA [11, 12]. Moreover, the studies focusing on circRNA-miRNA-mRNA networks have not developed a computational tool that automates the search for large-scale ce-circRNA [13, 14, 15, 16, 17, 18, 19]. Two circRNA functional annotation databases predicting sponge mechanisms involving circRNAs in several tissues have been developed, but they do not allow users to query their own transcriptomic datasets on a large scale, they most often limite queries to circRNA identifiers or sequences [20, 21]. Recently, a computational Nextflow pipeline has been developed that covers the circRNA and miRNA identification and quantification from sequencing data and includes a module for the search of ce-circRNA [22]. For the identification of ceRNA networks, the Nextflow pipeline uses the SPONGE R package [23], which was not initially designed to detect ce-circRNA as it requires miRNA and mRNA expression expression matrix as input. Although it presents an interesting approach, SPONGE provides ce-relationship between targets of miRNA but does not provide precise ce-mechanisms in the form of circRNA-miRNA-mRNA subnetworks. Finally, it is worth noting that, contrarily to the tool we propose, the SPONGE tool requires paired samples between each type of RNA expression data and is restricted to sequencing-based data within the Nextflow pipeline.

Here, we present Cirscan (CIRcular RNA Sponge CANdidates), an interactive Shiny application specifically developed to (i) infer circRNA-miRNA-mRNA networks based on human multi-level transcript expression data (circRNAs, mRNAs, and optionally miRNAs in the case of cancer study, with internal cancer-specific miRNA signatures) from two biological conditions (e.g. tumor versus normal conditions) provided by the user and by any technology (microarray or RNA-seq), and (ii) identify sponge mechanisms, active in a specific condition. The originality of our approach is to take into account multiple criteria (i.e. binding affinity, expression level) arising from ce-circRNA mechanisms to infer accurate ce-circRNA candidates on a large scale. In addition, our multiple criteria strategy leads to the calculation of a sponge score for each circRNA-miRNA-mRNA subnetwork, that greatly facilitates the prioritization of experimental validations.

We evaluated Cirscan on public multi-level transcript expression dataset from two cancers: colorectal cancer [24] and hepatocarcinoma cancer [25] and showed that Cirscan was able to retrieve previously described sponge mechanisms in the top ranked circRNA-miRNA-mRNA subnetworks, as well as identifying novel circRNA sponge candidates.

In summary, Cirscan provides a user-friendly tool to identify and visualize circRNA-miRNA-mRNA networks with potential sponge mechanisms from users human multi-level transcript expression data, and may provide novel therapeutic targets. Cirscan Shiny app is freely available on Gitlab (https://gitlab.com/geobioinfo/cirscan/_Rshiny). The scripts used in this paper are also provided on Gitlab (https://gitlab.com/geobioinfo/cirscan_paper).

## Implementation

### Overview of the tool

Cirscan infers circRNA-miRNA-mRNA interactions from coding and non-coding transcriptomic data (Fig. 1A) and identifies circRNA candidates that can act as miRNA sponges (ce-circRNA). The tool takes as input transcriptomic data from two biological conditions (e.g. tumor versus normal conditions), restricted to differentially expressed entities from any technology (microarray or RNA-seq). It requires from the user to normalized and log-transformed the data and a minimum of 3 samples per condition is required to ensure reliability of the downstream analysis. The tool comprises the following main steps: 1) Construction of a score matrix including multiple criteria based on interaction reliability and effectiveness of a sponge mechanism as defined in the following sections (Fig. 1B. Step 1), 2) Calculation of a sponge score for each circRNA-miRNA-mRNA subnetwork (Fig. 1B. Step 2). Cirscan outputs the top ranked circRNA sponge candidates based on the ranked sponge scores. It also enables the visualization of the identified sponge mechanisms as networks and biological enrichment analysis of the genes in the networks of interest (Fig. 1C). A video tutorial and an example dataset are available online to guide the user along the different steps of the application (https://gitlab.com/geobioinfo/cirscan_Rshiny).

**Fig. 1.**
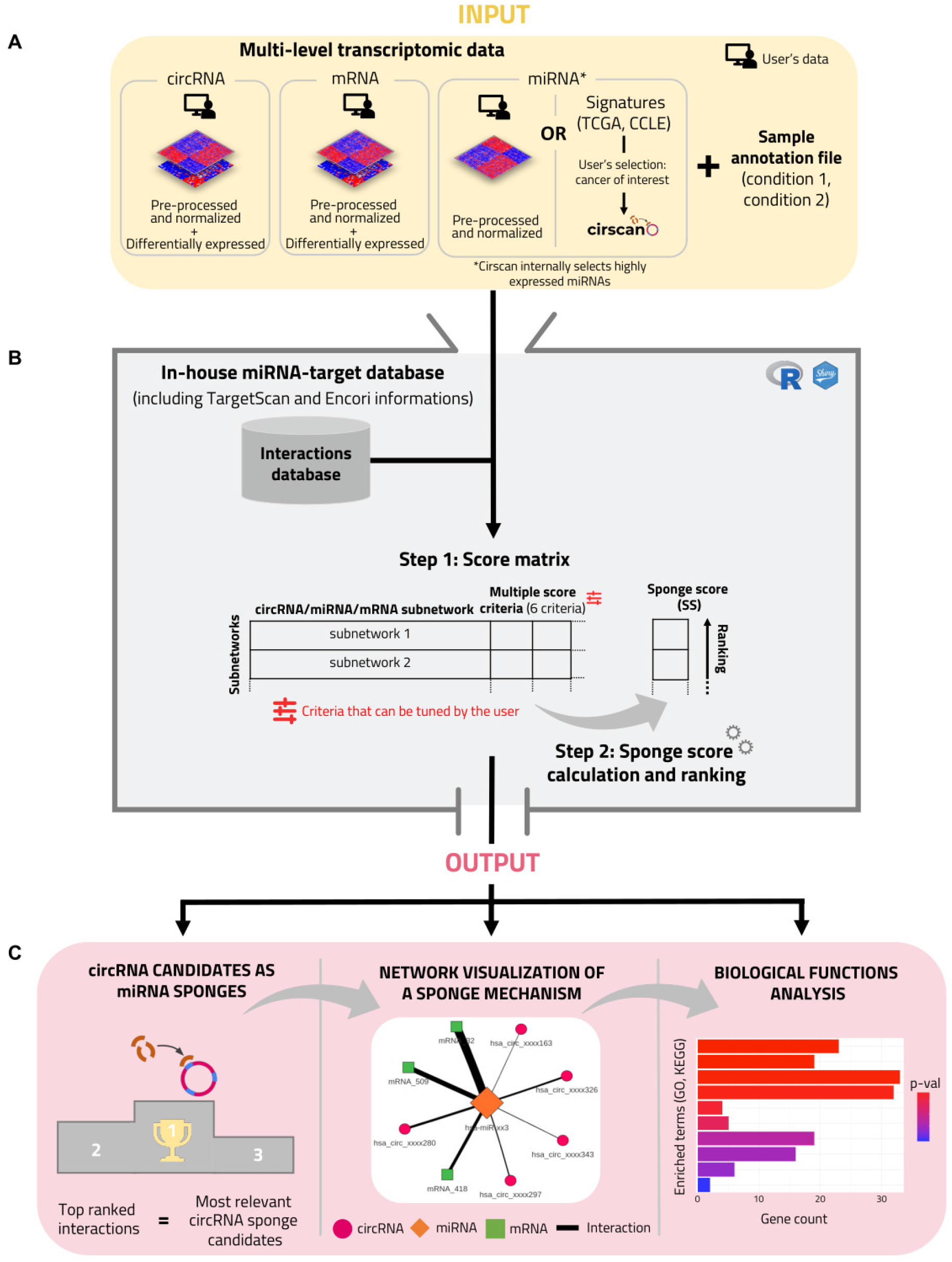
Cirscan workflow. (A) Input files (at least the circRNAs and mRNAs required) and a sample annotation file provided by the user. (B) Integration of multiple criteria for the calculation of a predicted sponge score for each circRNA-miRNA-mRNA subnetwork. (B. Step 1) Construction of the RNA score matrix based on interaction reliability and on RNA expression criteria. (B. Step 2) Calculation of a sponge score for each subnetwork. (C) Output: visualisation of the identified sponge mechanisms as networks around a selected RNA. The thickness of the edges is proportional to the sponge score and the RNA node of interest is bigger than the others. Additionally, a biological enrichment of mRNAs of networks of interest is performed.

### Effective sponge mechanisms prerequisites

The identification of circRNAs acting as miRNA sponges requires considering both sequence and expression information. Several prerequisites have been established to ensure the effectiveness of this identification process: 1) A miRNA must be sufficiently expressed whatever the biological conditions to have a functional impact on its targets and must target at least one circRNA and one mRNA, with a sufficient affinity. Importantly we do not assume a differential expression of the miRNA between the two conditions as we hypothesise that a sponge mechanism involving a circRNA affects the bioavailability of the miRNA but not its expression level. 2) Targets of the miRNA (mRNAs and circRNAs) must be co-expressed, i.e differentially expressed between the two conditions in the same direction. Indeed, it is expected that the increased presence of a circRNA that can sponge a miRNA will induce an over-expression of its mRNA targets, as the miRNA can no longer induce their degradation.

### Input files

Cirscan requires two types of CSV files from the user. First, multi-level transcript expression data are required as input files, including circRNA, mRNA and optionally miRNA expression matrices. If the user does not have miRNA expression data, miRNA signatures for different types of cancer (respectively 32 and 20 from the TCGA consortium (https://www.cancer.gov/tcga) and CCLE database (www.broadinstitute.org/ccle, [26])) are internally available in Cirscan. There is no constraint concerning the technology used to obtain the expression matrices (array or sequencing), which in return requires the user to properly normalize his data beforehand. Each expression matrix needs to be normalized (e.g. RMA for expression array, TPM for RNA-seq data) and log-transformed, and a minimum of 3 samples per condition is required to ensure reliability of the downstream analysis. Based on the sponge mechanism prerequisites, circRNA and mRNA expression matrices should be restricted to the differentially expressed circRNAs and mRNAs prior to the use of Cirscan. As mentioned above, this restriction is not relevant for miRNAs and no filter is required. Cirscan will automatically keep miRNAs that are sufficiently expressed considering the following assumptions: (1) the sum of the expression of a given miRNA in all samples must be nonzero, (2) for a given miRNA, its expression must be higher that a defined cutoff in more that 90% of the samples, the cutoff being the quartile Q1 of the overall miRNA expression distribution. This selection was also made for the establishment of the miRNA signatures from the TCGA (https://www.cancer.gov/tcga) and CCLE (www.broadinstitute.org/ccle, [26]) datasets.

CircRNAs identifiers must correspond to the human circRNAs referenced in circBase (“hsa circ xx xxxxx”) [27], human miRNAs from miRBase (“hsa-miR-xxx” or “hsa-let-xxx”) [28], and mRNAs must be referenced as official gene symbols. Furthermore, the user must provide another CSV file corresponding to the sample annotation file. This file must contain two columns: samples with the sample names referenced in the RNA expression matrices and conditions with the corresponding condition it refers to (condition 1, condition 2), knowing that the first condition refers to the control condition. The file format required by Cirscan is detailed in the Home Panel of the Shiny application and in the example dataset provided in the Interaction Ranking Panel.

### Identification of sponge mechanisms

#### In-house miRNA-target interaction database

A miRNA-target interaction occurs between the seed sequence of the miRNA and a complementary sequence called the miRNA Recognition Element (MRE) on the target (circRNA or mRNA). We used TargetScan [29] to predict miRNA-circRNA and miRNA-mRNA interactions, providing an affinity score for each interaction. For miRNA-circRNA interactions, we modified the affinity score calculated by TargetScan (context+ score, [30]) by excluding the 3’ pairing feature since it is not relevant for circRNA. To reduce false positive predictions, we set a cutoff for the context+ score based on its distribution in experimentally validated interactions extracted from the ENCORI database [31]. The cutoff was defined as the 95th percentile of the context+ score distribution for interactions supported by at least two CLIP-seq and degradome-seq experiments [see more details in Supplementary Material and Figure S1].

For miRNA-mRNA interactions, we used the affinity score provided by TargetScan that incorporates additional criteria relevant to miRNA-mRNA interactions that minimize false positive predictions, such as 3’ UTR length, ORF length, and probability of conserved targeting between species (context++ score, [29]) [see more details in Supplementary Material].

In total, the miRNA-target interaction database includes 1,244,932 miRNA-mRNA interactions and 31,045,182 miRNA-circRNA interactions.

#### Criteria of interaction reliability and sponge effectiveness

In order to identify potential sponge mechanisms involving circRNA, different scores of reliability are calculated. We defined the affinity score *S_Affinity_* between a miRNA and it targets as the context score calculated by Targetscan rescaled between 0 and 1: the closer to 1, the stronger the interaction affinity. For each miRNA-target interaction, we also considered the number of MRE sites that we named *S_NbMRE_*. If an interaction involves multiple MREs for the same miRNA and target of interest, the interaction with the maximum *S_Affinity_* is retained. To reduce the potential bias in the number of predicted MREs due to the length of the circRNA target sequence, the number of MREs for miRNA-circRNA interactions was normalized by the length of the circRNA sequence as described in the Cerina tool [21] and referenced here as the *S_NbMRE_*. In order to identify circRNAs that are significantly enriched for MRE binding sites of specific miRNAs, we used a binomial model conditioned on the total number of MRE for the given miRNA on each circRNA and the number of all other miRNA-MRE sites on the given circRNA as background, as proposed in the recent scanMiR tool [32]. The adjusted p-value (Benjamini-Hochberg, BH) of the enrichment test is defined as *S_EnrichMRE_*. A circRNA-miRNA interaction is kept if the Binomial enrichment test adjusted p-value (BH) is lower than 0.05.

In order to take into account the expression level information, different scores are calculated: the target expression fold change between conditions 1 and 2 (*S_FC_*), the median expression of each miRNA under all conditions (*S_MirExpr_*) and the highest mean expression of each target between the two conditions (*S_TargetExpr_*). Only networks with at least one circRNA and one mRNA with fold change values in the same direction are selected.

#### Calculation of a sponge score

For each miRNA-mRNA interaction, Cirscan calculates a global score defined as *SG_miRNA-mRNA_*, which integrates the different criteria described above *S_Affinity_*, *S_NbMRE_*, *S_FC_*, *S_MirExpr_*, *S_TargetExpr_* using the TOPSIS method [33]. Briefly, the TOPSIS method is a multi-criteria decision analysis method which aims to determine the best alternative when different criteria need to be considered together. In our case, the best alternative for each miRNA-mRNA interaction consists in maximizing the following criteria: highest affinity score (*S_Affinity_*), high number of MRE binding site (*S_NbMRE_*) and highly expressed miRNAs and differentially expressed circRNAs and mRNAs (*S_MirExpr_*, *S_TargetExpr_*, *S_FC_*).

Then, a score for each miRNA-circRNA interaction is calculated and defined as a sponge score, *SS*. The sponge score integrates the scores *S_Affinity_*, *S_NbMRE_*, *S_EnrichMRE_*, *S_FC_*, *S_MirExpr_*, *S_TargetExpr_*, *S_EnrichSG_*, where *S_EnrichSG_* corresponds to the adjusted p-value (BH) of the enrichment test of the miRNA-mRNA interactions for the miRNA of interest over all the miRNA-mRNA interactions ranked on the *SG_miRNA-mRNA_* score (R package fgsea). In other words, for a given circRNA-miRNA-mRNA subnetwork, if the miRNA-mRNA interactions have high *SG_miRNA-mRNA_* scores, this will contribute to increasing the sponge score. This sponge score is calculated using the TOPSIS method and in our case, the best alternative for each miRNA-circRNA interaction consists in maximizing the following criteria; *S_Affinity_*, *S_NbMRE_*, *S_MirExpr_*, *S_TargetExpr_*, *S_FC_*, and minimizing the adjusted p-values of enrichment tests, *S_EnrichSG_* and *S_EnrichMRE_*.

MiRNA-circRNA interactions are then ranked according to their sponge score, the top ranked interactions with the highest scores being the most relevant sponge circRNA candidates. Finally, all this information is compiled into a score matrix with 16 columns available and detailed in Cirscan, where *Rank_miRNA-circRNA_* is the rank of miRNA-circRNA interactions based on the sponge score *SS* and the top ones involving the most relevant sponge circRNA candidates.

### Visualization of the sponge mechanisms of interest

The identified circRNA-miRNA-mRNA subnetworks involving potential sponge mechanisms can be visualized in the Network Visualization Panel using the visNetwork R package (v2.0.8, https://github.com/datastorm-open/visNetwork). It is possible to visualize a network by selecting an interaction from the score matrix or by specifying the name of an RNA of interest: nodes correspond to the different types of RNAs (orange miRNAs, green mRNAs, and pink circRNAs) and edges to the miRNA-target interactions. The edge thickness is proportional to the sponge scores (the higher the value, the thicker the edge and the stronger the interaction affinity) and the RNA node of interest is larger than the others.

### GO and KEGG enrichment analysis of mRNAs

In order to help the user in the biological interpretation of the visualized network, an enrichment of biological terms on the genes of the visualized network is performed using Enrichr tool [34] with KEGG (KEGG 2021 Human) and GO (GO Biological Process 2021) databases.

## Results

### Cirscan performance evaluation

We evaluated Cirscan on two public multi-level transcript expression datasets from colorectal cancer (CRC) (n=20, 10 normal and 10 tumor samples) [24] and hepatocarcinoma (HCC) (n=14, 7 normal samples and 7 tumor samples) [35] in order to identify sponge mechanisms active in normal or tumor conditions.

For these two cancers, we retrieved sponge mechanisms referenced in the NcPath database [36], as well as described by Chen et al. [24] and in a recent review by [37]. We selected sponge mechanisms involving circRNAs for which a circBase identifier exists and a reference is associated, i.e. 30 sponge mechanisms described in the literature in colorectal cancer and 93 in hepatocarcinoma. The tables of sponge mechanisms are provided on our Gitlab web page (https://gitlab.com/geobioinfo/cirscan_paper).

Using the Cirscan tool on the CRC dataset pre-processed [see more details in Supplementary Material], we identified 12,850 potential circRNA-miRNA sponge mechanisms (i.e. 1,413 unique circRNAs) [Table available at https://gitlab.com/geobioinfo/cirscan paper as well as Supplementary File 1]. Among them, 3 sponge mechanisms were already described in the literature in CRC [38, 39] and significantly enriched in the top ranked sponge mechanisms given by Cirscan (GSEA enrichment [40], p-value = 4.04e-07) (Figure 2). Similar results were observed using the TCGA CRC miRNA signature available internally in Cirscan (p-value = 1.51e-06) [Table available at https://gitlab.com/geobioinfo/cirscan_paper as well as Supplementary File 2].

**Fig. 2.**
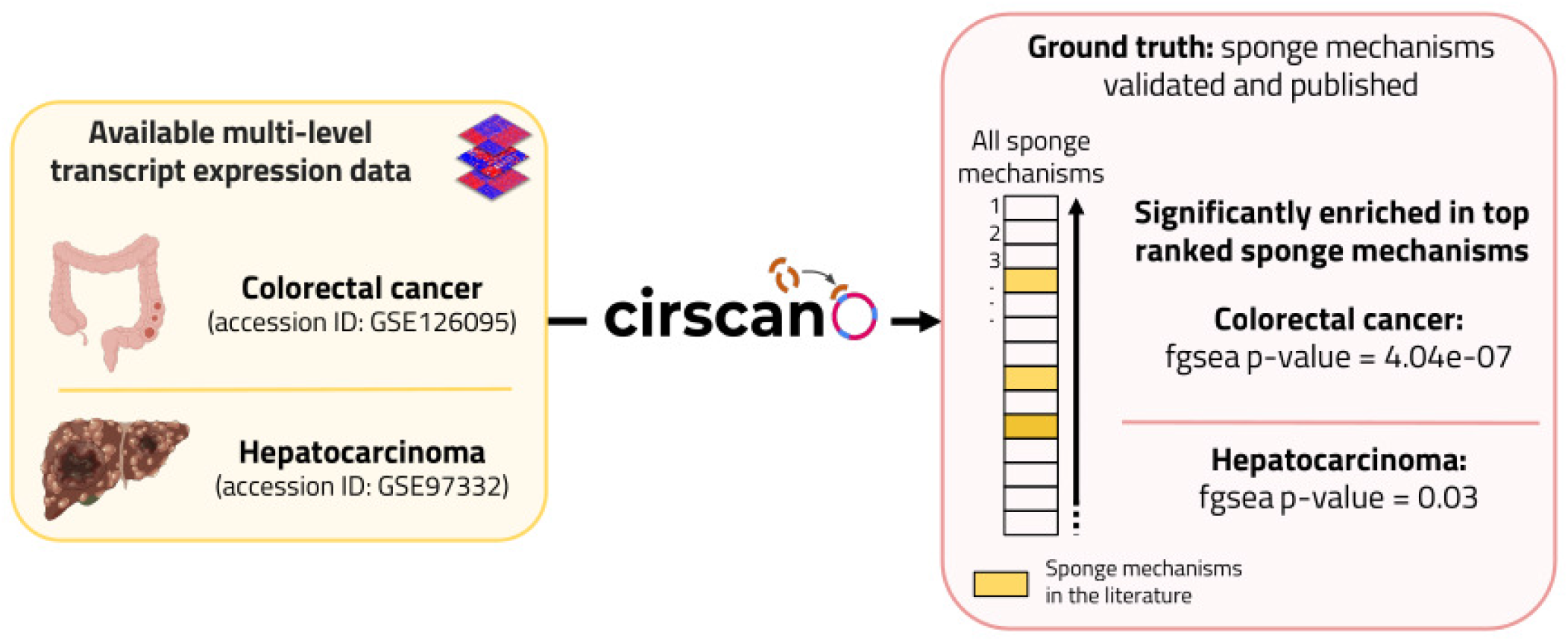
Cirscan performance evaluation. Significant enrichment of the sponge mechanisms identified in the literature in colorectal cancer and hepatocarcinoma data among the top candidates of Cirscan. Figure was created with BioRender.com.

The same approach was applied to the HCC circRNA and mRNA transcriptomic datasets and the TCGA HCC miRNA signature available internally in Cirscan, as no miRNA expression dataset was generated by the same authors [see more details in Supplementary Material]. We identified 16,190 potential circRNA-miRNA sponge mechanisms (i.e. 2,089 unique circRNAs) [Table available at https://gitlab.com/geobioinfo/cirscan_paper as well as Supplementary File 3] including 3 sponge mechanisms already described in the literature in HCC [41, 42, 43] that were also significantly enriched in the top ranked sponge mechanisms (p-value = 0.03) (Figure 2).

This significant enrichment of the sponge mechanisms identified in colorectal cancer and hepatocarcinoma in the literature among the top candidates of Cirscan confirms the reliability of our tool.

### Identification of known and novel sponge mechanisms in colorectal cancer

Using the public colorectal cancer dataset, Cirscan identified the hsa circ 0000977:hsamiR-135b-5p subnetwork, described in the literature by Ding et al. [38], at rank 18/12,850 out of all identified mechanisms (in the top 1%) and rank 17/5,468 out of all mechanisms identified in the normal condition (Figure 3.A). This result was consistent with the study of Ding et al. [38], as the authors describe this mechanisms as downregulated in the tumor condition. The KEGG biological term enrichment analysis of the mRNAs of this subnetwork revealed biological pathways associated with cell proliferation and differenciation involving the tumor suppressor genes APC and SMAD2 (Figure 3.B). Among the sponge mechanisms involved in the tumor condition and described in the literature, Cirscan identified the hsa circ 0001806:hsa-miR-193a5p subnetwork (Figure 3.C) (rank 105/12,850 (top 1%) and rank 32/7,382 (top 1%) mechanisms identified in the tumor condition) and the hsa circ 0001955:hsa-miR-145-5p subnetwork (rank 119/12,850 (top 1%) and rank 37/7,382 (top 1%)). The result of the KEGG enrichment of the hsa circ 0001806:hsa-miR-193a-5p subnetwork is consistent with the involvement of this sponge mechanism in the cancer with the enrichment of the TCF3 oncogene in the transcriptional misregulation in cancer pathway (Figure 3.D).

**Fig. 3.**
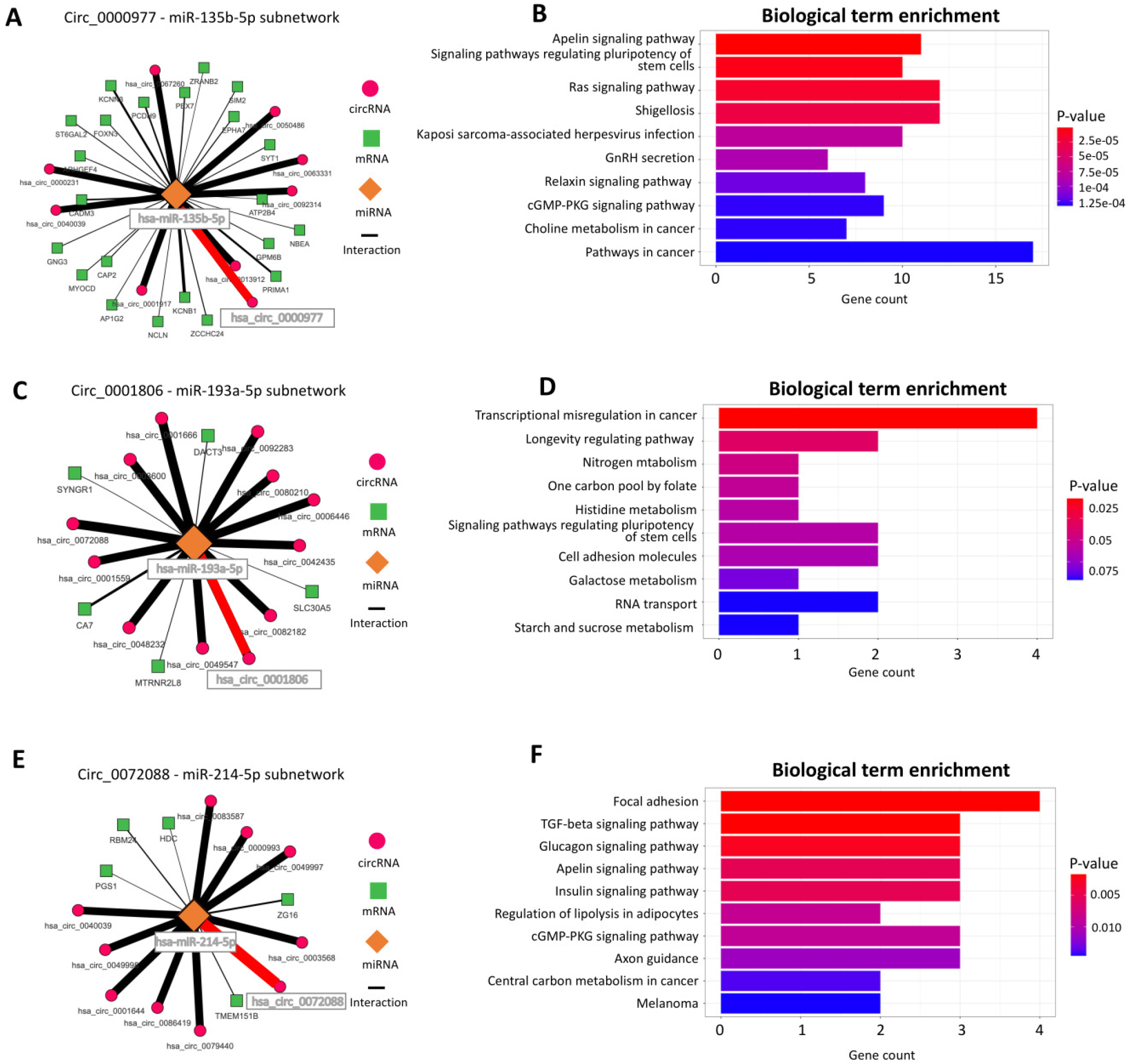
Subnetworks identified by Cirscan using colorectal cancer data. (A) Circ 0000977-miR-135b-5p subnetwork. The nodes represent the different RNA types (orange miRNAs, green mRNAs and pink circRNAs), and the edges represent miRNA-target interactions. The red edges represent the interactions found in the literature and the thickness of the edges is proportional to the interaction score values. For a better visibility, only the first 10% of the ranked targets are represented. (B) Top 10 mRNA-enriched pathways in the Circ 0000977-miR-135b-5p subnetwork using KEGG enrichment. (C) Circ 0001806-miR-193a-5p subnetwork, with the same legend as figure 3.A. (D) Top 10 mRNA-enriched pathways in the Circ 0001806-miR-193a-5p subnetwork using KEGG enrichment. (E) Circ 0072088-miR-214-5p subnetwork, with the same legend as figure 3.A. (F) Top 10 mRNA-enriched pathways in the Circ 0072088-miR-214-5p subnetwork using KEGG enrichment.

We next focused on the best ce-circRNA candidates having a sponge role in the tumor condition. Interestingly, within the top 10 ce-circRNA candidates, the first ce-circRNA has been already shown to have a sponge role in lung and colorectal cancer in several studies (hsa circ 0072088: [44, 45, 46, 47]). Cirscan highlighted potential novel sponge mechanisms for this circRNA, involving for example the miRNA hsa-miR-214-5p (Figure 3.E) targeting genes associated to signaling pathways enriched with oncogenes, such as PDGFRA and AKT3 (Figure 3.F). Other ce-circRNAs within the top 10 ce-circRNA candidates in the tumor condition have been identified in other cancers as hepatocellular carcinoma, lung or esophageal cancer (hsa circ 0000517 [48, 49], hsa circ 0000326 [50], hsa circ 0000518 [51]). The other best ce-circRNA candidates (hsa circ 0000644, hsa circ 0011385, hsa circ 0006174, hsa circ 0003945, hsa circ 0008274, hsa circ 0043438) have not yet been described in the literature as associated with cancer, making them interesting subjects for further experimental validations.

### Identification of known and novel sponge mechanisms in hepatocarcinoma

The same approach was applied to the HCC transcriptomic data. Cirscan identify the hsa circ 0074854:hsa-miR-338-3p subnetwork, ranked 1,279/16,190 out of all identified mechanisms (in the top 10%) and ranked 639/8,468 out of all mechanisms identified in the tumor condition (Figure 4.A), as well as described by Li et al. [42]. The biological term enrichment analysis of the mRNAs of this subnetwork revealed pathways associated with cancer, such as the enrichment of the GNAQ, STAT1 and AKT3 oncogenes in the MAPK signaling pathway (Figure 4.B). Moreover, Cirscan identified at lower ranks the subnetworks hsa circ 0000130:hsa-miR-141-3p (rank 2,764/16,190 and rank 1,430/8,468) and hsa circ 0001461:hsa-miR-30a-5p (rank 14,037/16,190 and rank 7,333/8,468) already described in the literature ([41] and [43]), making them relevant mechanisms in the study of HCC.

**Fig. 4.**
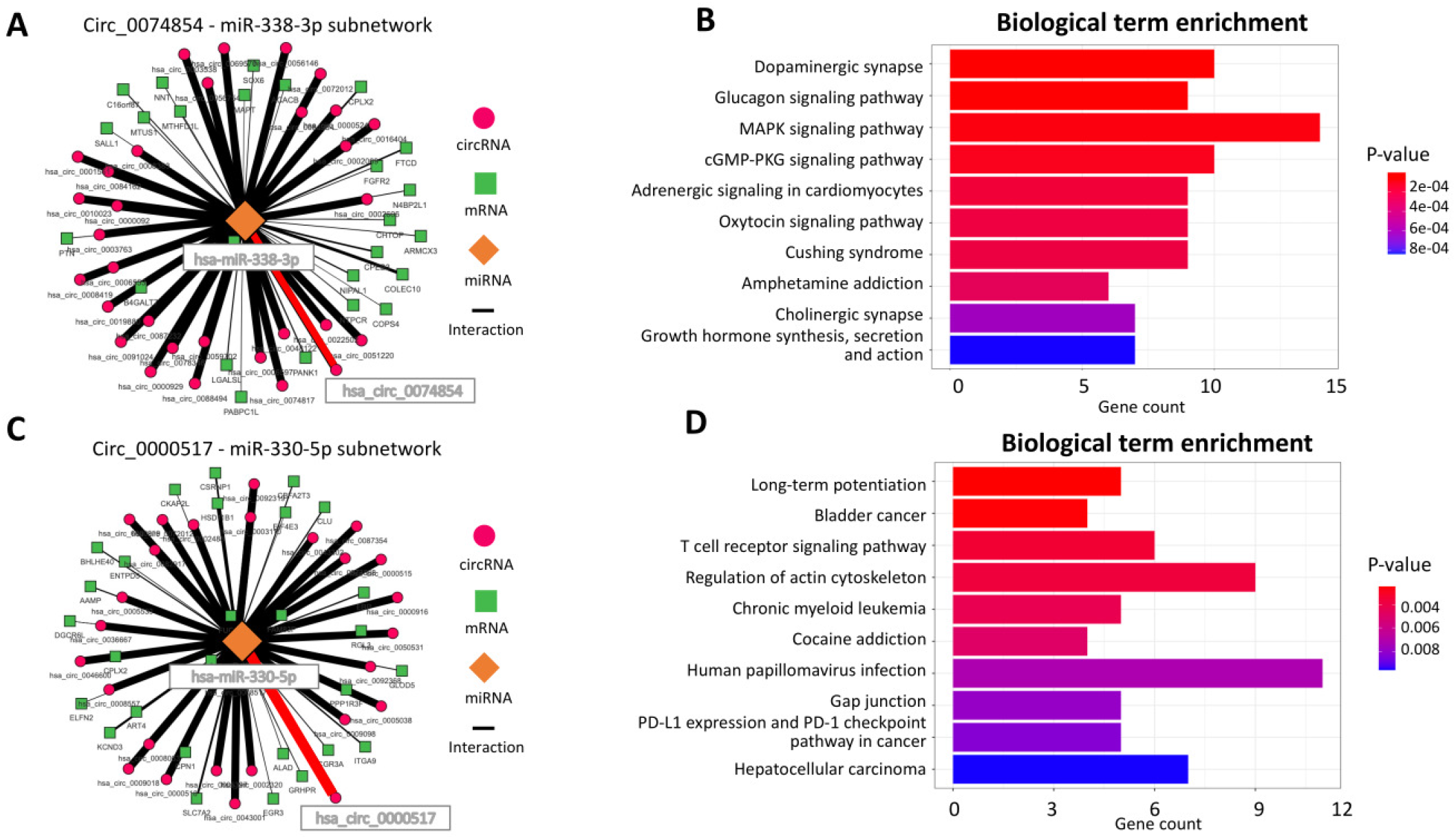
Subnetworks identified by Cirscan using hepatocellular cancer data. (A) Circ 0074854-miR-338-3p subnetwork. The nodes represent the different RNA types (orange miRNAs, green mRNAs and pink circRNAs), and the edges represent miRNA-target interactions. The red edges represent the interactions found in the literature and the thickness of the edges is proportional to the interaction score values. For a better visibility, only the first 10% of the ranked targets are represented. (B) Top 10 mRNA-enriched pathways in the Circ 0074854-miR-338-3p subnetwork using KEGG enrichment. (C) Circ 0000517-miR-330-5p subnetwork, with the same legend as figure 4.A. (D) Top 10 mRNA-enriched pathways in the Circ 0000517-miR-330-5p subnetwork using KEGG enrichment.

Interestingly, within the top 10 ce-circRNA candidates described by Cirscan and active in the tumor condition, the third hsa circ 0000517 (Figure 4.C) and the seventh hsa circ 0001834 were already described in the literature as differentially expressed in HCC [48, 52]. Cirscan identified a potential novel sponge mechanisms for the hsa circ 0000517, involving the hsa-miR-330-5p that targets genes, for example the oncogene NRAS, enriched in different cancers, such as hepatocellular carcinoma and bladder cancer (Figure 4.D). Among the 10 ce-circRNA candidates, hsa circ 0048122 is identified as being involved in colorectal cancer [25] and hsa circ 0000644 in bladder cancer [53]. The other best ce-circRNA candidates (hsa circ 0000516, hsa circ 0087643, hsa circ 0052523, hsa circ 0046599, hsa circ 0042521, hsa circ 0064288) have not yet been described in the literature as associated with cancer, making them interisting subjects for further experimental validations.

## Conclusions

Cirscan is a tool that takes as input any human multi-level transcript expression data to identify on a large scale and visualize condition-specific sponge mechanisms involving circRNAs. As shown on two different applications, Cirscan allows the identification of known and novel potential sponge mechanisms that may be further investigated and validated experimentally. This tool can be considered as a companion tool for biologists, facilitating their ability to prioritize sponge mechanisms for experimental validations. The mechanisms revealed by Cirscan could open new avenues for the development of novel RNA-targeted therapies. In particular, it would be possible to use antisense oligonucleotides, which can bind by complementarity to circRNA sequences of interest to inhibit the sponge mechanisms active in a specific condition [54]. Finally, the framework established in this study could be extended to other species.

## Availability and requirements

Project name: Cirscan

Project home page: https://gitlab.com/geobioinfo/cirscan_Rshiny

Project application page: https://gitlab.com/geobioinfo/cirscan_paper

Operating system(s): Platform independent

Programming language: R and CSS Other requirements: R 4.2.2

Licence: GNU General Public Licence Restrictions for academic use: none

## List of abbreviations

BH: Benjamini-Hochberg
ce-circRNA: Competitive Endogenous circRNA
circRNA: Circular RNA
Cirscan: CIRcular RNA Sponge CANdidates
CRC: Colorectal Cancer
HCC: Hepatocarcinoma
MRE: miRNA Recognition Elements
miRNA: microRNA
ncRNA: Non-coding RNA

## Declarations

### Ethics approval and consent to participate

Not applicable

### Consent for publication

Not applicable

### Availability of data and materials

Cirscan is implemented in R, released under the license GPL-3 and accessible on GitLab (https://gitlab.com/geobioinfo/cirscan_Rshiny). The scripts used in this paper are also provided on Gitlab (https://gitlab.com/geobioinfo/cirscan_paper).

### Competing interests

The authors declare that they have no competing interests.

### Funding

This study received financial support from the Ligue National Contre le Cancer (LNCC) Dpartements du Grand-Ouest. RMF is a recipient of a doctoral fellowship from the LNCC and YSA is a recipient of a doctoral fellowship from the LNCC Grand Ouest.

### Authors’ contributions

YB supervised the project and conceptualized the tool. YB & RMF developed and implemented the Cirscan tool. MDG, SC & YSA contributed to the biological inputs. MA contributed to the in-house interaction database construction. YB & RMF wrote the original draft of the manuscript. All authors read, improved and approved the final manuscript.

## Acknowledgments

The authors would like to thank the Gene Expression and Oncogenesis team (UMR6290) for helpful discussions. We also acknowledge the GenOuest bioinformatics core facility for providing the computing infrastructure and BioRender software (BioRender.com) for the creation of Figure 2.

## Notes

https://gitlab.com/geobioinfo/cirscan_Rshiny

https://gitlab.com/geobioinfo/cirscan_paper

### Competing Interest Statement

The authors have declared no competing interest.

