## supplementary_material for "Cirscan: a shiny application to identify differentially active sponge mechanisms and visualize circRNA-miRNA-mRNA networks"

**3 Supplemental References** **3**

### 1 Cirscan Description

#### 1.1 In-house miRNA-target interaction database

A miRNA-target interaction is established between a region of the miRNA, called "seed sequence" and a complementarity sequence called "miRNA Recognition Element" (MRE) on the target (circRNA or mRNA). An in-house database of interactions between miRNAs and putative mRNA or circRNA targets was constructed using the TargetScan interaction prediction tool (McGeary et al. 2019) and information from experimentally validated interaction database (ENCORI, (Li et al. 2014)).

TargetScan was used to predict miRNA-circRNA and miRNA-mRNA interactions, and to obtain an affinity score for each interaction, by taking into account the specificity of the RNA type considered (mRNA or circRNA). For miRNA-circRNA interactions, we used 140,790 circRNA sequences from circBase (Glažar, Papavasileiou, and Rajewsky 2014) and 9,994 miRNA sequences from TargetScan (Garcia et al. 2011). TargetScan v6 calculates for each miRNA-circRNA interaction an affinity score called context+ score, corresponding to the sum of the contribution of six features (site-type, 3' pairing, local AU, position, target site abundance and seed-pairing stability). The feature 3' pairing was removed here because it is not relevant for circRNA as the pairing is not expected to be restricted to that region and can appear on the whole circRNA sequence. The lower the context+ score, the stronger the affinity between the miRNA and its target. To reduce the number of false positive predictions, we defined a cutoff on the context+ score, based on its distribution on experimentally

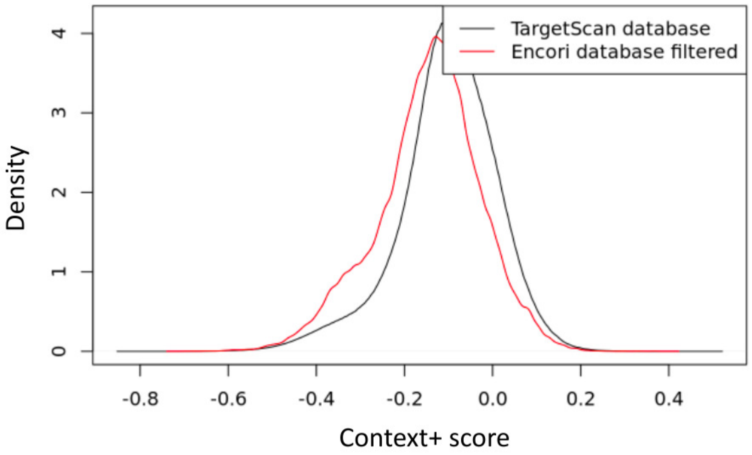

Figure S1: Context+ score distribution on miRNA-circRNA interactions predicted by the TargetScan database and on experimentally validated miRNA-circRNA interactions (CLIP-seq and degradome-seq >= 2) by the ENCORI database.

validated interactions given by the ENCORI database. We defined as a cutoff the 95th percentile of the context+ score distribution restricted to the interactions supported by at least two CLIP-seq and degradome-seq experiments (Figure S1). We used the ENCORI/Starbase API [`http://starbase.sysu.edu.cn/tutorialAPI.php`] to download the interactions contained in this database.

We retrieved 2,055,407 miRNA-circRNA interactions:

```
curl 'https://starbase.sysu.edu.cn/api/miRNATarget/?assembly=hg19&
    geneType=circRNA&miRNA=all&clipExpNum=0&degraExpNum=0&pancancerNum=0&
    programNum=1&program=None&target=all&cellType=all'
```

and 1,215,274 miRNA-mRNA interactions:

```
curl 'https://starbase.sysu.edu.cn/api/miRNATarget/?assembly=hg19&
    geneType=mRNA&miRNA=all&clipExpNum=0&degraExpNum=0&pancancerNum=0&
    programNum=1&program=None&target=all&cellType=all'
```

For miRNA-mRNA interactions, we used 2,382,569 UTR sequences and 9,994 miRNA sequences from TargetScan (Garcia et al. 2011). TargetScan v8 calculates for each miRNA-mRNA interaction an affinity score called context++ score (Agarwal et al. 2015). Compared to the context+ score, the context++ score includes additional criteria specifically relevant for miRNA-mRNA interactions (e.g. 3' UTR length, ORF length, probability of conserved targeting between species), which are expected to reduce false positive predictions.

MiRNA-circRNA and miRNA-mRNA interactions were restricted to specific MRE binding sites: 7mer-m8 (exact match to positions 2-8 of the mature miRNA), 7mer-1a (exact match to positions 2-7 of the mature miRNA followed by an "A"), and 8mer-1a (exact match to positions 2-8 of the mature miRNA followed by an "A"), i.e. 1,244,932 miRNA-mRNA interactions and 31,045,182 miRNA-circRNA interactions in total.

### 2 Cirscan performace evaluation

#### 2.1 Pre-processing of colorectal cancer data

Microarray multi-level transcript expression data of 10 colorectal cancer (CRC) samples and 10 normal adjacent samples were downloaded from the NCBI Gene Expression Omnibus database (accession number: GSE126095). Each dataset was imported into the RStudio (v1.4.1103) environment with R (v4.2.2) for pre-processing.

For transcripts belonging to the gene annotation, an expression average was applied. Quantile normalization and a log-transformation were applied to the mRNAs expression matrix. mRNAs microarray matrix was reduced to protein-coding genes, using the annotation file provided by the authors. Using the limma R package (v.3.50.1) (Ritchie et al. 2015), 4,640 differentially expressed mRNAs with an adjusted p-value (Benjamini-Hochberg, BH) $< 0.05$ were selected. The circRNAs microarray matrix was also submitted to quantile normalization and a log-transformation. We selected 1,491 differentially expressed circRNAs with an adjusted p-value (BH) $< 0.05$ by using the limma R package (v.3.50.1) (Ritchie et al. 2015). Finally, the miRNAs microarray matrix was filtered to only include miRNA identifiers present in the annotation file, i.e. 2,055 miRNAs. Quantile normalization and log-transformation were also applied to the miRNAs expression matrix. These different pre-processed expression matrices were given as input to Cirscan.

#### 2.2 Pre-processing of hepatocarcinoma data

circRNAs microarray expression data of 7 hepatocellular carcinoma (HCC) tissues and 7 non-tumor liver tissues were downloaded from the NCBI Gene Expression Omnibus database (accession number: GSE97332). mRNAs expression matrix of 424 liver samples (374 tumor tissues and 50 normal tissues) were downloaded from the TCGA (`https://www.cancer.gov/tcga`). Each dataset was imported into the RStudio (v1.4.1103) environment with R (v4.2.2) for pre-processing.

circRNAs identifiers have been converted according to the nomenclature described by circBase (i.e. 3,416 unique circRNAs). The circRNAs microarray expression matrix was restricted to 2,222 differentially expressed circRNAs with an adjusted p-value (BH) $< 0.05$ by using the limma R package (v.3.50.1) (Ritchie et al. 2015). mRNAs counts expression matrix was reduced to protein-coding genes, using the annotation file provided by using the Ensembl database and the Biomart web tool (Howe et al. 2021, `https://www.ensembl.org/index.html`). As mRNA expression data was already log-transformed, we performed a conversion to raw data ($2 \wedge$x-1 transformation) in order to use them with the DESeq2 R package (v1.40.2) (Love, Huber, and Anders 2014), and we selected 10,760 mRNAs differentially expressed with an adjusted p-value (BH) $< 0.05$. These different pre-processed expression matrices were given as input to Cirscan. Finally, the miRNAs expression data was directly retrieved from the signature of sufficiently expressed miRNA from liver cancer available in the Cirscan tool, as no miRNA expression dataset was generated by the same authors.
